# Chemodiversity defines a fundamental synecological dimension of plant form and function

**DOI:** 10.64898/2026.07.22.740092

**Authors:** Maximilian Hanusch, Alexander Zizka, Robert R. Junker

## Abstract

Plant trait spaces have advanced ecology by reducing the vast diversity of plant form and function to a few axes of trait variation. Yet, such frameworks remain dominated by autoecological traits related to resource acquisition and growth, while largely overlooking the traits mediating interactions with other organisms. Chemodiversity, the richness, evenness and chemical disparity of volatile organic compounds (VOCs) emitted by leaves and flowers to attract, deter, or otherwise affect biotic interaction partners, may represent such an overlooked synecological dimension of plant functional variation. By integrating the chemodiversity of floral (*n* = 859 species) and vegetative (*n* = 159 species) VOC profiles into the global spectrum of plant form and function, we show that chemodiversity is a) independent from the classical axis of variation in plant size and leaf economics but b) is linked to biotic interactions in organ-specific ways. Flower-visitor richness increases with floral scent chemodiversity, a pattern supported by an analysis of global interaction data and a meta-analysis of the specialization and generalization of flower-animal interactions. Vegetative VOC chemodiversity has context-dependent positive and negative effects on herbivore richness. VOC chemodiversity represents a distinct ecological strategy that defines interaction niches, thereby contributing to ecological differentiation and coexistence among otherwise functionally similar species.

## Introduction

Plant traits reflect the ecological strategies that underlie the diversity of plant form and function (Carmona and Beccari 2025). Trait spaces condense this diversity into a few meaningful dimensions of functional and ecological variation. Traditionally, these dimensions have rested on autecological traits that capture fundamental trade-offs in growth and resource-use strategies, such as the spectra of plant size and leaf economics (Díaz et al. 2016; Dehling and Stouffer 2018). More recently, trait space approaches have incorporated traits that additionally reflect the synecology of plants, i.e. their interactions with their biotic environment, by adding morphological traits of flowers or roots as further dimensions (Bergmann et al. 2020; E-Vojtkó et al. 2022). As these traits capture distinct aspects of plant function, they carry largely non-redundant information and form axes decoupled from the classical economics spectra (Carmona et al. 2021). So far, however, this expansion has relied mostly on morphology, even though many plant biotic interactions are mediated by chemical information (Zimmer 2000; Vlot and Rosenkranz 2022; Hanusch et al. 2026). Adding a chemical dimension would therefore capture aspects of a plant’s ecological strategy that cannot be adequately represented by morphology alone.

A plant’s chemical phenotype is a key aspect of its interaction niche (Müller and Junker 2022; Chadwick et al. 2026) and plants produce a high diversity of secondary (specialized) compounds to interact with their physical and biological environment (Bouwmeester et al. 2019). In principle, each metabolite could be represented as a separate dimension within a trait space. However, this would provide little informative gain, while making the data increasingly sparse and obscuring meaningful patterns in high-dimensional space (Bellman 1966). Rather than accounting for each metabolite individually, aggregate indices describing properties of metabolomes or volatilomes condense their compositional information and thereby offer a tractable alternative (Walker et al. 2023; Yang et al. 2026). Accordingly, we propose chemodiversity, an index that captures the richness, evenness, and biochemical disparity of blends of metabolites (Petrén et al. 2023), as a dimension in trait spaces as an index that meets all criteria of a functional trait sensu Violle *et al*. (Violle et al. 2007): Chemodiversity is a physiological trait that is measurable at the level of an individual, and affects the individual’s fitness via effects on survival and reproduction (Müller et al. 2025).

Like specific leaf area, a cornerstone of plant trait spaces, chemodiversity is a derived measure associated with many plant functions and physiological processes (Wetzel and Whitehead 2020). While chemodiversity is a relatively recent concept, its ecological functionality is empirically supported, with demonstrated effects on herbivores, pollinators, and microbes (Richards et al. 2016; Petrén et al. 2024; Hanusch et al. 2025). However, when chemodiversity is quantified from untargeted metabolomics spanning primary and secondary metabolites, contributions from functionally unrelated compounds potentially obscure its association with any specific ecological strategy and thereby limit its informative value as a trait space dimension. To avoid the functional overdispersion captured in metabolomes, volatile organic compound (VOC) emissions from leaves and flowers may provide a more targeted chemical measure, as they are largely composed of compounds with demonstrated functions in biotic interactions (Dötterl and Gershenzon 2023). Additionally, the biosynthetic pathways of VOCs released by plants are often well known facilitating the consideration of biochemical disparity in chemodiversity measures (Junker 2018; Petrén et al. 2023).

While vegetative VOCs contribute to plant tolerance of abiotic stress (Zuo et al. 2025), their functions mostly comprise biotic interactions ranging from the repellence of herbivores and the attraction of herbivores’ enemies (even beyond the third trophic level; Poelman et al., 2008) to plant-plant communication (Erb 2019), thereby strongly contributing to the survival and growth of plants. VOCs emitted by flowers are mainly associated to plant reproduction by repelling floral antagonists and attracting pollinators (Junker and Blüthgen 2010; Dötterl and Gershenzon 2023). VOCs emitted by leaves and flowers are also involved in shaping the microbial communities associated with aboveground plant parts (Junker and Tholl 2013; Hammerbacher et al. 2019; Adler et al. 2021). All of these functions have mostly been attributed to individual VOCs, which raises the question why leaf and flower scent bouquets are composed of multiple compounds (Junker 2016). A number of evolutionary hypotheses have been proposed to explain the chemodiversity of plants (Wetzel and Whitehead 2020; Thon et al. 2024) that can be summarized as “the more, the better”, which means more compounds can fulfill more functions and mediate interactions with a diverse set of organisms (Thon et al. 2024). Empirical studies support this notion (Petrén et al. 2024) and suggest that chemodiversity mediates the generalization / specialization of interaction networks (Larue et al. 2016; Hanusch et al. 2025). Chemodiversity therefore constitutes a synecological aspect of a plant’s phenotype that promotes ecological differentiation among co-occurring species with otherwise similar autecological strategies, thereby facilitating their coexistence (Schiestl and Johnson 2013; Sedio et al. 2017).

In this study, we extracted data on the VOC emission of flowers (*n* = 859 plant species) and leaves (*n* = 159) from the literature and quantified the chemodiversity of these bouquets using the R package *chemodiv* (Petrén et al. 2023). We then included the chemodiversity of plant-species and -organ specific VOC emissions into the classical set of traits used to define the global spectrum of plant form and function (Díaz et al., 2016, 2022). To confirm the notion that chemodiversity is a functional trait *sensu* Violle et al. (2007), we tested whether it is related to the number of animal species interacting with flowers and leaves. As chemodiversity is expected to mediate biotic interactions, it may be distinct from the classical traits that mediate autecological functions, such as stem specific density or leaf nitrogen content. We therefore hypothesized chemodiversity to represent an independent dimension of the global spectrum of plant form and function. Alternatively, chemodiversity can be hypothesized to be constrained by plant morphology and physiology, which would lead to correlations between chemodiversity and the major axes of the global spectrum of plant form and function. Resolving between these alternatives would determine whether VOC chemodiversity is a redundant echo of plant morphology and physiology, or a distinct, synecological dimension of plant ecological strategy with implications for community assembly and the coexistence of otherwise functionally similar species.

## Methods

### Compilation of plant volatile profiles

We compiled data on VOCs emitted by flowers and leaves through a systematic review of the existing literature. Relevant studies were identified by a search in the Web of Science Core Collection on April 23, 2024. We used two separate search strings for studies reporting either floral or vegetative volatile profiles. For floral profiles, we used the following search string: (flower OR floral) AND (head$space OR volatile* OR scent OR scent emission OR volatile organic compound* OR odo$r) AND (composition OR bouquet OR profil*), which yielded *n* = 2,343 records. For vegetative volatiles, we applied a similar search string: (leaf OR leaves) AND (head$space OR volatile OR scent OR scent emission OR volatile organic compound OR odo$r) AND (composition OR bouquet OR profil*), resulting in *n* = 2,987 records. In both cases, the search was limited to studies published from 1980 onwards.

All records and corresponding abstracts were downloaded and screened using the R package metagear (v0.7, Lajeunesse, 2016). Abstracts were evaluated for suitability based on the following predefined criteria: We excluded studies focusing on hybrids, including only wild or parental (non-hybrid) individuals. Similarly, ornamental breeds or energy plants were excluded, and crop species were screened individually, retaining those that retain naturally evolved, ecologically functional interactions (e.g. animal pollination and intact floral emission). Furthermore, we retained only studies that analyzed spontaneously emitted volatiles, i.e. compounds naturally released from plant tissues under ambient or controlled conditions. Studies relying on solvent extraction, maceration, or other invasive techniques were excluded because these methods do not reflect constitutive emission profiles. Finally, we included only studies that analyzed headspace samples using gas chromatography coupled with mass spectrometry to ensure methodological comparability in compound detection and identification across studies.

After abstract screening, we retained *n* = 469 studies on floral scent and *n* = 227 studies on vegetative volatiles (Fig. S1). We extracted data on volatile emissions from these studies if i) the full text was accessible, ii) concentrations of individual compounds were reported, and not mere presence/absence data only, and iii) compound abundance was quantified using mass-based, molar, concentration-based, or signal-based measures, including peak areas or intensities. Based on these criteria, we were able to collect data from *n* = 247 floral scent and *n* = 73 vegetative volatile studies. Data was extracted either manually or using the software Tabula v.1.2.1(Aristarán et al. 2018) from tables in the main text or from supplementary data of individual studies.

### Data curation and standardization

Following the compilation of raw data, we performed a series of steps to standardize the records for subsequent quantitative analyses.

First, all compound records with missing, zero, or ambiguous abundance values that indicate their absence (e.g. “– “, “NA”, or “0”) were removed. Entries indicating trace amounts, such as values denoted by symbols (e.g. “<”, “!”, “tr”, “nd”), were assigned a small non-zero value of 0.01 irrespective of its unit to conservatively acknowledge their presence allowing inclusion in quantitative analyses.

Second, chemical compound names were standardized to IUPAC chemical nomenclature to remove inconsistencies across studies and annotated with chemical identifiers (canonical SMILES and InChIKeys). Identifiers were retrieved from PubChem using either CAS numbers from the original studies or, when unavailable, the compound names provided in the study. Automated queries were performed in R using the packages httr v.1.4.7 (Wickham 2023) and *jsonlite* v.2.0.0 (Ooms 2014). Records that could not be resolved through automated queries were manually reviewed. In total, we were able to assign standardized chemical identifiers to 86.6% of all recorded compounds, i.e. *n* = 41,914 of 48,429 compounds.

Third, plant taxon names extracted from the literature were standardized to currently accepted nomenclature by resolving spelling inconsistencies and taxonomic synonyms. This was achieved by matching plant taxon names against the World Flora Online Taxonomic Backbone using the R package *WorldFlora* v1.14-5 (Kindt 2020).

Fourth, we screened the dataset for duplicate records of floral profiles to avoid repeated measurements originating from the same sampling event. We defined duplicates as entries reporting the same compounds with identical abundance values for the same plant species. Such duplicates can arise when chemical profiles of individual plants are reused across multiple publications, for example through shared datasets or reanalysis of previously published data. When potential duplicates were detected, we consulted the corresponding publications to evaluate whether the entries originated from the same underlying dataset or represented independent measurements. Based on this screening, we identified duplicate records originating from nine studies in the floral scent dataset and one study in the vegetative volatile dataset and removed the corresponding entries.

After all data curation steps, the final dataset comprised *n* = 39,781 individual compound measurements across *n* = 859 plant species for floral scent, and *n* = 8,648 compound measurements across *n* = 159 plant species for vegetative volatiles.

### Calculation of Chemodiversity

Chemodiversity was calculated using the R package chemodiv v.0.3.0 (Petrén et al. 2023), which is specifically designed to quantify the chemical diversity of complex mixtures such as floral scents or vegetative volatile bouquets. We calculated functional scent diversity from pairwise structural dissimilarities among compounds based on PubChem fingerprints. PubChem fingerprints represent each compound as a binary vector of predefined substructure features, and pairwise dissimilarity was calculated from fingerprint similarity scores of these substructures. For compounds that remained unidentified, we assigned them the overall mean dissimilarity value. We used the resulting dissimilarity matrix to calculate functional Hill diversity, a metric that quantifies the effective total abundance weighted dissimilarity between compounds within a sample (Chiu and Chao 2014), with the sensitivity parameter set to *q* = 1, a commonly used value at which the measure is sensitive to compound richness, evenness, and dissimilarity, and therefore the most comprehensive measure of chemodiversity (Petrén et al. 2023). Chemodiversity was calculated at the individual sample level and subsequently aggregated at the species level by averaging chemodiversity values to their mean across all sampled individuals per species. Interspecific variability exceeded intraspecific variability in compound richness (mean absolute difference between species: 6.68; within species: 3.67; 1.82-fold), as well as in functional Hill diversity (q=1; between species: 27.0; within species: 17.6; 1.54-fold), indicating that individuals of the same species are chemically more similar to one another than to individuals of other species. The resulting chemodiversity-by-species matrix formed the basis for all subsequent analyses and is hereafter referred to as either the *floral* or *vegetative chemodiversity dataset*.

### Testing the functionality of chemodiversity

To confirm that chemodiversity constitutes a trait that is functional, we investigated its role in shaping the degree of generalization versus specialization in plant–animal interactions. Specifically, we focused on mutualistic interactions with pollinators and antagonistic interactions with herbivores integrating two complementary approaches: First, we complemented our chemodiversity data with estimates of interaction partner richness from the Global Biotic Interactions database (GloBI; Poelen et al., 2014) to capture large-scale patterns in realized interaction networks across a broad taxonomic and geographic scope. Second, we conducted a meta-analysis of studies quantifying either floral scent or vegetative chemodiversity and their relationship with interaction partner richness. The meta-analysis allowed us to assess the strength and consistency of these relationships under more controlled and well-characterized study conditions.

### Statistical analysis of chemodiversity–species richness relationships

To test whether chemodiversity predicts the number of interaction partners at a broad taxonomic and geographic scale, we related chemodiversity to interaction partner richness extracted from GloBI. We accessed the GloBI database via the R package rglobi v. 0.3.4 (Poelen et al. 2026) on March 24, 2025. For each plant species in either the floral or vegetative chemodiversity datasets, we queried interaction records with either the interaction type “pollinatedBy” (for flowers) or “eatenBy” (for leaves). We conservatively cleaned the resulting datasets from obviously implausible entries. Implausible entries include, for instance, self-links where the plant itself is listed as an interaction partner, as well as associated taxa that are neither pollinators nor above-ground herbivores (e.g., bacteria, fish, or earth worm); these were removed at the family level (Supporting Information 2B). We found interaction partner entries for *n* = 322 species and *n* = 78 species of the species according to the floral and the vegetative chemodiversity dataset.

We tested the relationship between chemodiversity and interaction partner richness while accounting for phylogenetic non-independence among species using a species-level phylogeny based on a global megatree (https://raw.githubusercontent.com/megatrees/plant_20221214/main/GBOTB.extended.WP.tre; Jin and Qian 2022; Smith and Brown 2018; Zanne et al. 2014) using the R packages ape v. 5.8-1 (Paradis and Schliep 2019), U.PhyloMaker v.0.1.0 (Jin and Qian 2023) and nlme v.3.1-168 (Pinheiro et al. 2025). The tree was pruned to include only species present in the chemodiversity dataset; two species (*Prosopanche americana* and *P. bonacinae*) could not be matched and were excluded from phylogenetically informed analyses. We then fitted phylogenetic generalized least squares models, incorporating a Pagel’s λ correlation structure estimated by maximum likelihood during model fitting. Chemodiversity (measured as functional Hill diversity with *q* = 1) and interaction partner richness were log-transformed prior to analysis. Results were qualitatively unchanged when chemodiversity was quantified as chemical compound richness (*q =* 0).

### Meta-analysis of chemodiversity–species richness relationships

To test whether chemodiversity–richness relationships hold under more controlled and well-characterized study conditions than the large-scale GloBI data allow, we conducted a meta-analysis of studies quantifying chemodiversity in relation to interaction partner richness. The meta-analysis focused on two interaction contexts: i) chemodiversity of floral scent in relation to the diversity of flower-visitor assemblages, and ii) chemodiversity of vegetative tissues in relation to herbivore diversity. In general, literature reporting both chemodiversity and pollinator or herbivore richness remains sparse. Our synthesis thus represents the best available quantitative summary and is seen to complement the broader interaction-database analysis.

For the compilation of studies investigating floral scent chemodiversity, we relied on three studies identified during the literature research to collate the chemodiversity dataset, which already involved a comprehensive and systematic screening of floral scent literature. For vegetative chemodiversity, we draw on a compilation by Petren et al. (2023) that is based on a systematic literature search and synthesizes empirical studies explicitly linking plant chemical traits to metrics of herbivore community diversity. From this source, we extracted all studies (*n* = 45) that quantified vegetative tissue chemodiversity and reported corresponding measures of herbivore species richness or diversity. Eligible studies quantified a component of chemical α-diversity (e.g., compound richness, diversity, or functional diversity) and reported either extractable raw data or standardized regression coefficients. Based on these criteria, three studies on floral scent (five effect sizes) and six studies from the herbivore literature (24 effect sizes) were included. Given the limited number of eligible studies quantifying vegetative VOC and herbivore diversities, we decided to also include studies that characterized chemodiversity based on non-volatile metabolites.

Where available, standardized regression coefficients were extracted directly from the published studies. If only raw data or graphical outputs were provided, standardized coefficients were retrieved by manually extracting data points using WebPlotDigitizer (automeris.io). All analyses were conducted as multilevel random-effects meta-analyses using restricted maximum likelihood as implemented in the R-package metafor v4.8-0 (Viechtbauer 2025). To account for non-independence among multiple effect sizes derived from the same study, all models included random intercepts for study identity, with effect sizes nested within studies to account for non-independence. Potential small-study effects and publication bias were evaluated using funnel plots and formally tested with Egger’s regression.

### Global spectrum of plant form and function

To assess whether chemodiversity represents an independent axis of trait variation or is captured by the established global spectrum of plant form and function, we obtained a published, species-level plant trait dataset comprising species-mean values for six key functional traits that define the primary axes of covariation in this spectrum (Sandra Díaz et al. 2022). The trait dataset contained measurements of leaf area, leaf mass per area, plant height, stem specific density, leaf nitrogen content, and diaspore mass. It represents a partial trait matrix, with 36.86% of species-by-trait combinations missing. To obtain a complete dataset suitable for subsequent analyses, we imputed missing entries using Bayesian hierarchical probabilistic matrix factorization (BHPMF; Fazayeli et al., 2014), which is a taxonomically constrained gap-filling approach (Díaz et al. 2016; Bruelheide et al. 2018; Walker et al. 2023). Prior to imputation, traits were log-transformed and standardized, and a taxonomic hierarchy comprising species, genus, and family was supplied to inform the hierarchical imputation procedure following the approach of Schrodt et al. (2015). Cross-validation indicated good imputation accuracy (RMSE = 0.79). We screened the resulting dataset for outliers that reflect either measurement errors or imputation artefacts by manually inspecting all species with a standardized trait value (z-score) exceeding |4| for any of the trait values (i.e., deviating by more than 4 SD from the mean; Kattge et al. 2011; Díaz et al. 2016; Walker et al. 2023), which resulted in the removal of 9 species (i.e. < 2% of the total dataset; Supporting Information 2A). The imputed trait data were then merged with the chemodiversity dataset at the species level. In total, we yielded a complete dataset comprising a full matrix of the six functional traits and corresponding chemodiversity estimates for *n* = 350 plant species based on floral scent profiles, and *n* = 107 plant species based on vegetative volatile profiles.

### Chemodiversity as a distinct trait-space dimension

We assessed whether chemodiversity covaries with the major axes of the global spectrum of plant form and function or represents a separate dimension of trait variation using principal component analyses (PCA), conducted separately for the floral and vegetative datasets. Prior to analysis, chemodiversity was log-transformed and all variables were standardized to zero mean and unit variance. PCA was performed on the correlation matrix using the R package FactoMineR v2.1.3 (Lê et al. 2008). We examined the proportion of variance explained by each component and the variable loadings to evaluate the contribution of chemodiversity to the major axes of trait variation, and visualized relationships among all variables in three-dimensional PCA space.

### Independence of chemodiversity from major trait axes

We complemented PCA with an exploratory factor analysis (EFA) to identify underlying latent constructs of coordinated trait variation, allowing us to test whether chemodiversity is integrated into or independent of the major trait syndromes forming the global spectrum of plant form and function. Analyses were conducted using the R package psych (v. 2.5.6; William Revelle, 2026). Prior to analysis, chemodiversity was log-transformed and all variables were standardized to zero mean and unit variance. The suitability of the data for factor analysis was confirmed using Bartlett’s test of sphericity (floral: χ²(21) = 890.57, p < 0.001; vegetative: χ²(21) = 281.48, p < 0.001). The number of factors was determined by parallel analysis, which compares the eigenvalues from the observed data with those obtained from randomly generated datasets of the same size and supported the retention of two factors in both datasets. Factor extraction was performed using maximum likelihood estimation, and solutions were rotated using promax rotation to allow for correlated latent factors. To assess the robustness of factor loadings, bootstrapped confidence intervals were estimated based on 999 iterations. Factor loadings were examined to evaluate whether chemodiversity covaried with classical functional traits.

## Results

### Chemodiversity across species and organs

Floral scent chemodiversity (disparity-weighted functional Hill diversity, *q* = 1) averaged at 56 ± 78 (mean ± SD); the corresponding unweighted effective number of compounds (Hill number, *q* = 1) averaged at ∼7. The most complex bouquets occurred in groups such as the Brassicaceae and Orchidaceae with up to ∼26 effective compounds (functional Hill diversity ∼360). The least diverse scents, by contrast, belonged to the trap-flowered *Aristolochia microstoma* and hawkmoth-pollinated *Mandevilla* species (∼2 compounds; functional Hill diversity ∼4–8).

In the vegetative dataset, chemodiversity was on average higher than in the floral dataset (74.5 ± 100; ∼9 effective compounds). As our chemodiversity measure captures the richness, evenness, and chemical disparity of a blend rather than its perceived intensity, strongly aromatic but individual compound-dominated species, such as cineole-rich *Eucalyptus* species, ranked among the lowest (∼2–4 compounds; functional Hill diversity ∼2–11), whereas the most diverse blends occurred in *Populus* (∼26 compounds; functional Hill diversity ∼660) that combines many volatiles of comparable abundance and high chemical disparity.

### Testing the functionality of chemodiversity

Floral scent chemodiversity showed a positive effect on pollinator species richness (β = 0.16 ± 0.067 SE, *p* = 0.016; Fig. 1A). Vegetative volatile chemodiversity showed a non-significant negative relationship with herbivore species richness β = -0.07 ± 0.174 SE, *p* = 0.667; Fig. 1B).

**Figure 1:**
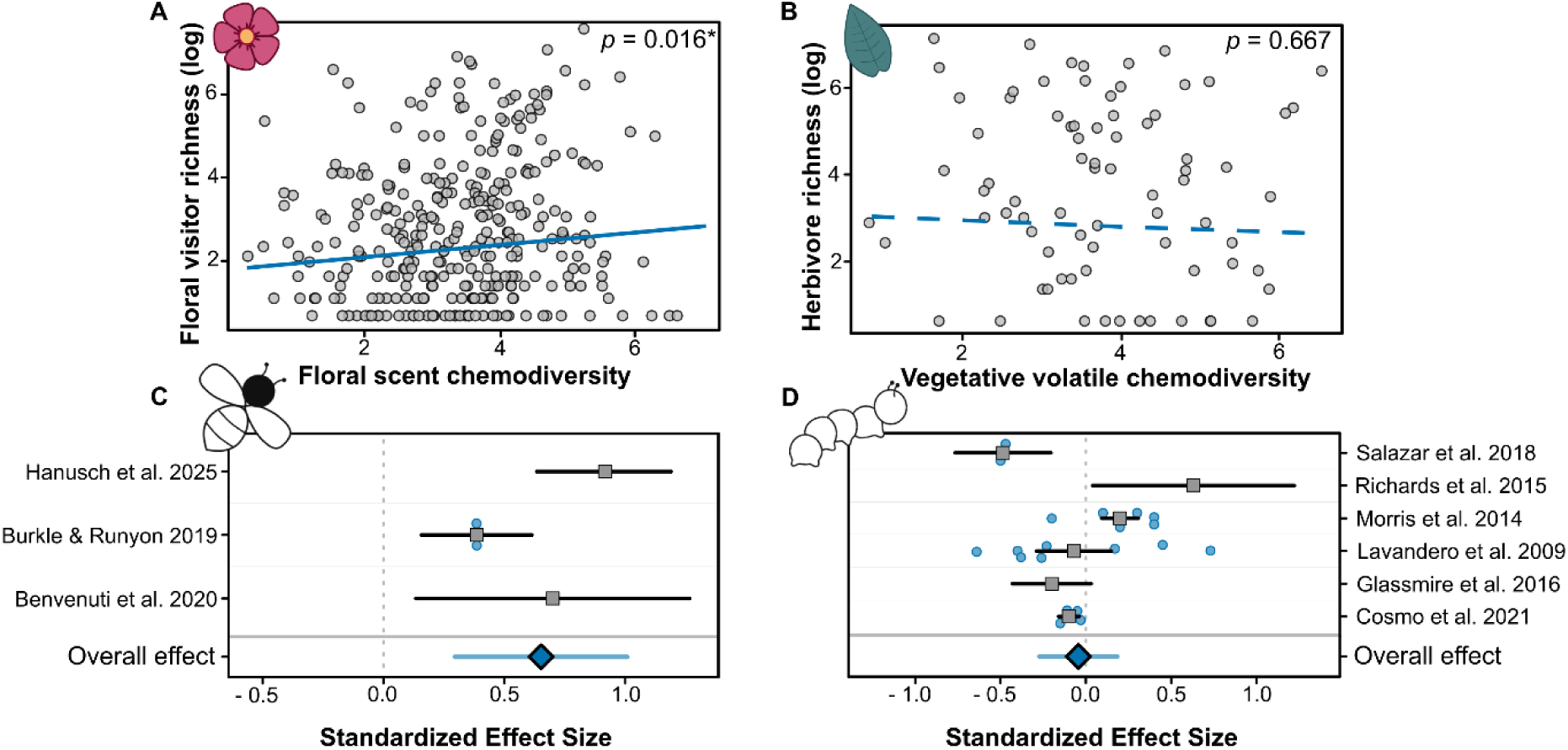
Chemodiversity of flowers and leaves and its relationship with diversity of interacting pollinators and herbivores. Top row: analyses based on the Global Biotic Interactions (GloBI) database, using phylogenetically informed models. (A) Floral scent chemodiversity is positively associated with pollinator richness across *n* = 322 plant taxa. (B) In contrast, vegetative scent chemodiversity shows no significant relationship with herbivore richness across *n* = 78 plant taxa. Bottom row: meta-analytic synthesis of published evidence. (C) Across 3 studies (5 effect sizes), floral scent chemodiversity shows a consistent positive association with pollinator richness. (D) Across 6 studies (24 effect sizes), vegetative chemodiversity shows no consistent overall relationship with herbivore richness, despite significant effects in both directions among individual studies.

The meta-analyses revealed contrasting patterns for floral and vegetative chemodiversity in structuring biotic interactions. For floral scent chemodiversity, we found a consistent positive association with floral visitor richness across three studies, supporting the pattern observed in the interaction database. The overall standardized effect size was significantly positive (estimate = 0.652, SE = 0.182, 95% CI = 0.296–1.008; z = 3.59, p < 0.001). The 95% prediction interval was also positive (0.030–1.275), showing that the positive association is robust across studies. In contrast, the meta-analysis of vegetative chemodiversity revealed no consistent association with herbivore richness. The overall standardized effect size was close to zero and not significant (estimate = −0.044, SE = 0.116, 95% CI = −0.272–0.183; z = −0.38, p = 0.70). However, effect sizes were highly heterogeneous (Q₂₁ = 72.30, p < 0.001), and the 95% prediction interval included both negative and positive effects (−0.625 – 0.537), indicating strong context dependence across studies.

### Chemodiversity as a distinct trait-space dimension

Principal component analysis revealed highly similar patterns for floral and vegetative chemodiversity (Fig. 2), characterized by a clear separation between the dominant axes of classical trait variation (PC1–PC2) and a distinct third axis (PC3) reflecting variation in chemodiversity. In both datasets, PC1 and PC2 largely reproduced the size and leaf-economics axes of the global spectrum of plant form and function (Díaz et al. 2016; Sandra Díaz et al. 2022) while chemodiversity instead defined an almost orthogonal third axis.

**Figure 2:**
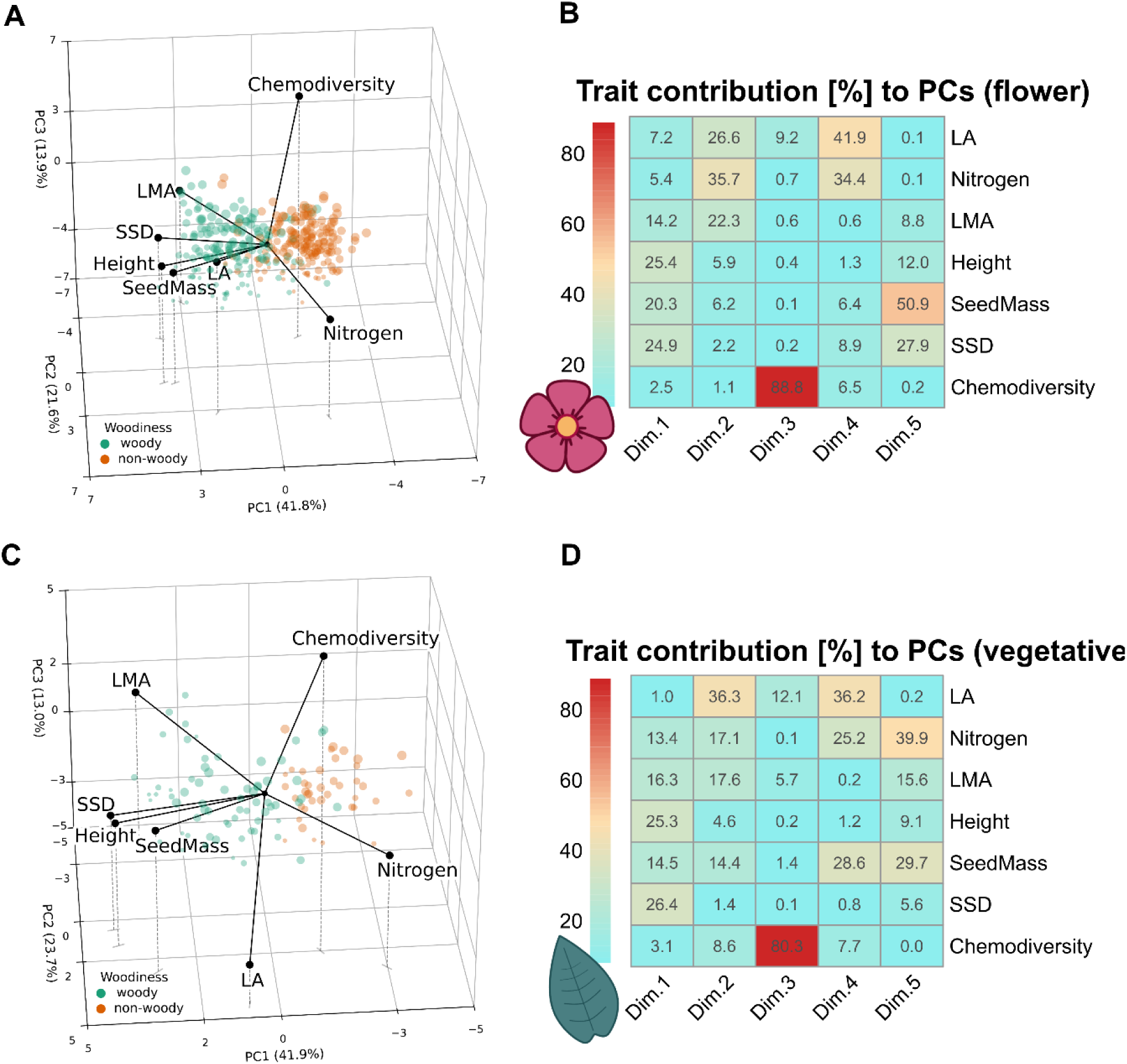
Principal component analysis (PCA) of floral and vegetative scent chemodiversity in relation to six classical functional traits representing major axes of plant form and function. Point size reflects chemodiversity (small = low, large = high), and point color indicates woodiness (green = woody, orange = non-woody species). Solid lines represent trait vectors (variable loadings), indicating the direction and strength of trait associations in multivariate space, while dashed lines aid visual interpretation. Variable contributions to individual principal components are visualized as a heatmap in which coloration reflects the contribution of each trait to each principal component (blue = low, red = high) (A) Three-dimensional PCA scores (PC1–PC3) for floral scent data and the six functional traits representing key axes of plant form and function (*n* = 351 species), showing that chemodiversity is largely orthogonal to the dominant trait variation captured by PC1 and PC2. (B) Variable contributions to the principal components for floral scent, indicating that chemodiversity contributes little to PC1 and PC2 but primarily and almost exclusively to PC3. (C) Three-dimensional PCA scores (PC1–PC3) for vegetative scent data (*n* = 107 species), showing a similar pattern of separation from the main trait axes. (D) Variable contributions to the principal components for vegetative scent, again indicating minimal contribution of chemodiversity to PC1 and PC2 and an almost exclusive association with PC3.

In the floral chemodiversity dataset, PC1 and PC2 explained 41.8% and 21.6% of total trait variation, respectively (63.4% cumulative), and PC3 explained an additional 13.9% (77.3% cumulative; Fig. 2A). PC1 was dominated by classical functional traits: plant height (25.4%), stem specific density (24.9%), and diaspore mass (20.3%), consistent with a size-related dimension, while PC2 was dominated by leaf nitrogen content (35.7%), leaf area (26.6%), and leaf mass per area (22.3%), consistent with a leaf-economics dimension. Floral scent chemodiversity contributed little to either axis (2.5% and 1.1%, respectively) but accounted for 88.8% of the variable contribution to PC3 (Fig. 2B).

In the vegetative chemodiversity dataset, PC1 and PC2 explained 41.9% and 23.7% of total variation (65.6% cumulative), and PC3 explained an additional 13.0% (78.6% cumulative). As in the floral dataset, PC1 was dominated by stem specific density (26.4%), plant height (25.3%), and diaspore mass (14.5%), while PC2 was dominated by leaf area (36.3%), leaf mass per area (17.6%), and leaf nitrogen content (17.1%), consistent with size and leaf-economics dimensions, respectively. Vegetative chemodiversity showed only weak contributions to these dimensions (3.1% and 8.6%; Fig. 2C) but instead contributed 80.3% to PC3 (Fig. 2D).

### Independence of chemodiversity from major trait axes

For the flower and vegetative chemodiversity datasets, exploratory factor analysis consistently identified two latent dimensions corresponding to the two major axes of plant trait variation identified in the global spectrum of plant form and function. In both analyses, one factor represented a plant size axis and the other a leaf economics axis, while chemodiversity did not load substantially on either factor (Fig. 3).

**Figure 3:**
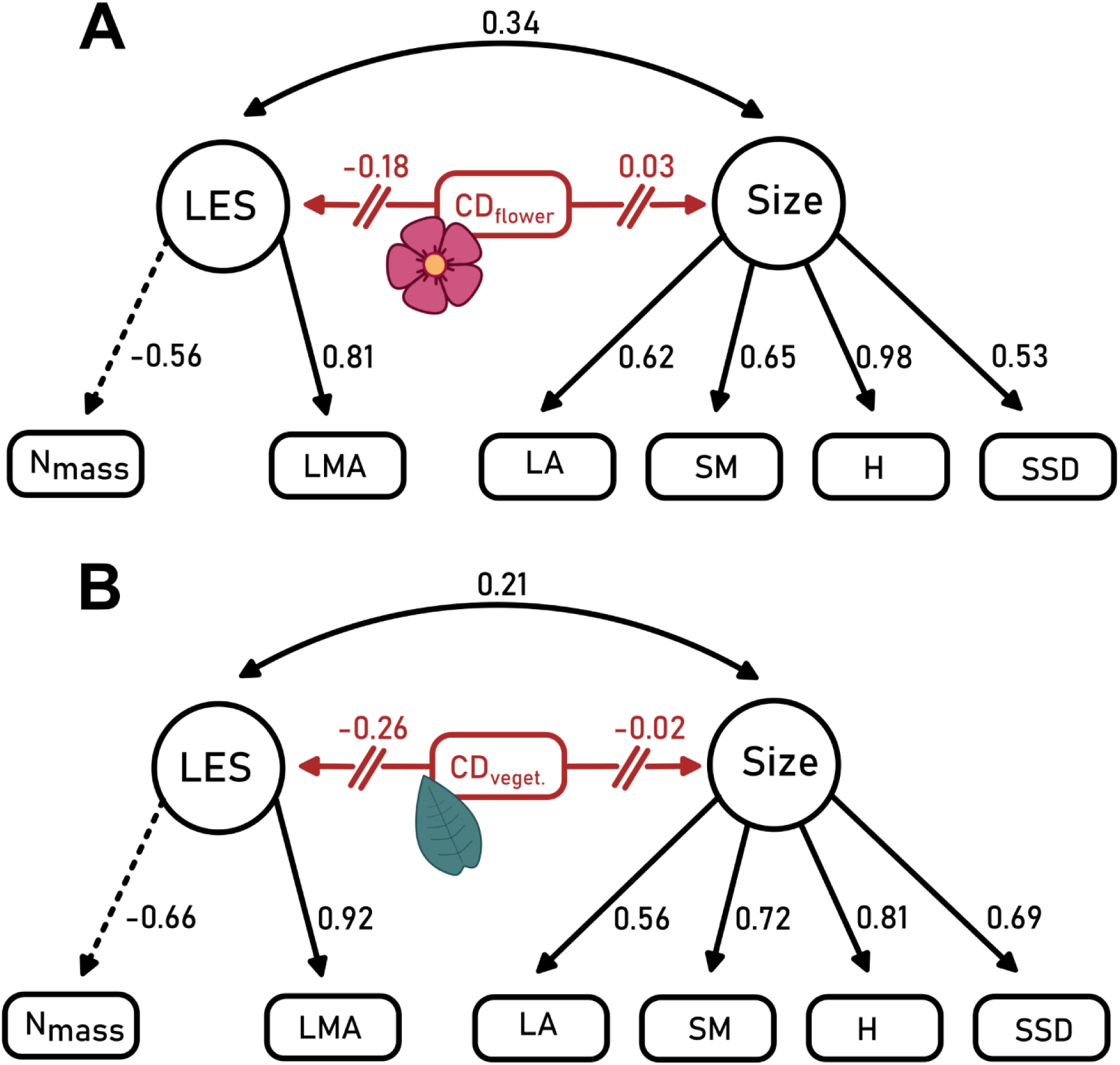
Exploratory factor analysis of floral scent and vegetative VOC chemodiversity and six functional traits representing key axes of plant form and function, showing a two-factor solution corresponding to a plant size dimension and a leaf economic dimension. Standardized factor loadings are indicated by arrows; the red crossed arrow for chemodiversity indicates negligible loadings on both factors (|loading| < 0.3). (A) Floral scent chemodiversity exhibits high uniqueness (u² = 0.96), indicating that its variance is largely not explained by the latent trait dimensions. (B) Vegetative chemodiversity shows weak loadings on both factors and high uniqueness (u² = 0.93), indicating limited integration into the dominant axes of trait variation.

In the flower dataset, the two-factor solution explained 52% of the total variance, with 31% attributable to the size-related factor and 21% to the leaf economics factor (Fig. 3A), under good model fit (TLI = 0.93, RMSEA = 0.09; Supporting Information 3). Bootstrapped confidence intervals for chemodiversity loadings overlapped zero for both factors, and uniqueness was high (u² = 0.96), indicating that 96% of the variance in floral scent chemodiversity is not captured by the two latent factors.

In the leaf dataset, the two-factor solution explained 55% of the total variance, with 29% attributable to the leaf economics factor and 26% to the size-related factor (Fig. 3B, under good model fit (TLI = 0.99, RMSEA = 0.03; Supporting Information 3). Bootstrapped confidence intervals overlapped zero for the size axis and ranged from −0.49 to −0.04 for the leaf economics axis; vegetative volatile chemodiversity also showed high uniqueness (u² = 0.93).

## Discussion

While our analyses draw on heterogeneous data sources, spanning multiple study systems, and levels of sampling effort, a consistent global pattern emerges: chemodiversity is largely independent from the two main axes of the global spectrum of plant form and function. Further, our results position the chemodiversity of floral and vegetative VOCs as a functional trait that complements the autecological strategies captured by the global spectrum of plant form and function with an independent synecological dimension. As such, chemodiversity contributes to shaping the degree of specialization *versus* generalization in plant biotic interactions. Across complementary approaches, a large-scale analysis of interaction records from the Global Biotic Interactions database and a meta-analysis of published studies, higher floral scent chemodiversity was consistently associated with higher flower-visitor richness. In contrast, vegetative VOC chemodiversity showed no consistent overall relationship with herbivore richness. Instead, associations varied in strength and direction among studies, suggesting strong context dependence.

The classical axes of plant size and resource economics primarily summarize autecological strategies of resource acquisition and allocation (Wright et al. 2004; Díaz et al. 2016). Chemodiversity is largely independent from these axes, indicating that it captures an additional, predominantly synecological dimension of plant strategy: although secondary metabolites and volatiles also mediate responses to abiotic conditions, much of their functional diversity reflects how a plant interacts with its associated biota rather than how it acquires and allocates resources (Kessler and Kalske 2018). Accordingly, species with similar growth and resource-use strategies can nonetheless differ substantially in their chemical profiles (Eilers et al. 2021; Müller and Junker 2022; Müller et al. 2026). Such chemical differentiation could allow plants to maintain distinct sets of interaction partners, thereby separating their interaction niches despite facing comparable ecological constraints and trade-offs (Gargallo-Garriga et al. 2020). This interaction niche separation represents a recognized route to coexistence among otherwise functionally similar species (Peeters et al. 2001; Marks and Lechowicz 2006; Becerra 2015; Dias et al. 2020).

The different patterns between floral and vegetative scent chemodiversity could also partly reflect fundamental differences in the regulation of organ-specific VOC emissions. Floral emissions are largely constitutive (Raguso 2008), whereas vegetative VOCs are predominantly inducible and dynamically regulated in response to herbivory (Kessler and Baldwin 2001). Consequently, the literature-derived measurements of vegetative chemodiversity included in our analysis may not capture the realized defensive phenotype expressed under herbivore pressure. In addition, the ecological effects of vegetative volatile and non-volatile chemodiversity depend on herbivore feeding strategies (Volf et al. 2015b, a). Higher chemodiversity may suppress generalist herbivores through additive and synergistic effects between compounds, whereas specialist herbivores may be adapted to specific host-associated compounds, which they tolerate, detoxify, or sequester, and may therefore be less affected or even favored (Ali and Agrawal 2012; Richards et al. 2016). Accordingly, plant species specific differences in whether their associated herbivore assemblages are dominated by generalists or specialists can lead to relationships between chemodiversity and herbivore richness that vary in their magnitude and direction across systems (Lewinsohn et al. 2005). Moreover, vegetative chemodiversity is shaped by selective pressures unrelated to herbivory, including abiotic stress mitigation (Loreto and Schnitzler 2010). This context dependence likely obscures a single global relationship between chemodiversity and herbivore richness, consistent with our meta-analysis and previous studies reporting positive (e.g. Morris et al. 2014; Richards et al. 2015), as well as negative associations with herbivore richness (e.g. Salazar et al. 2018; Cosmo et al. 2021). Thus, rather than indicating a lack of function, this variability more likely reflects the context-dependent and multifunctional roles of vegetative chemodiversity.

We used flower-visitor attraction and herbivore repellence as tractable, well-established proxies to show the effects of volatile chemodiversity on the degree of specialization versus generalization of a plant species’ biotic interactions. However, evidence from other systems shows that the functionality of chemodiversity extends beyond these interaction types by influencing the diversity of interacting organisms across multiple trophic groups. For example, floral scent chemodiversity defines not only pollinator assemblages but also the diversity of floral microbial communities (Hanusch et al. 2025), while vegetative VOC chemodiversity has been shown to attract more diverse assemblages of parasitoids that act as natural enemies of herbivores (Wan et al. 2017). Thus, chemodiversity structures a plant’s degree of specialization versus generalization across a broader range of organismal groups, positioning it as a multifunctional plant trait with far-reaching consequences for plant fitness (Müller and Junker 2022).

Both floral scent and vegetative volatile chemodiversity were independent of the two major axes of the global spectrum of plant form and function. Thus, chemodiversity emerges as a distinct and ecologically meaningful dimension of plant trait variation, capturing biotic-interaction strategies that are not represented by the classical traits of plant size and resource economics. There is growing consensus that plant trait frameworks remain incomplete without explicit representation of traits that mediate biotic interactions (E-Vojtkó et al. 2022; Carmona and Beccari 2025; de Bello et al. 2025). Current descriptions of plant form and function have been recognized to be mainly centered around Grinnellian niche dimensions, i.e. the abiotic conditions under which a species persists, while giving less attention to Eltonian niche dimension, i.e. how plants interact with other organisms (Dehling and Stouffer 2018). Our study presents multiple independent lines of evidence indicating that chemodiversity is a functional trait relevant to biotic interactions and therefore captures information about Eltonian niche dimensions that is otherwise poorly represented by classical approaches. As such, chemodiversity might provide one of the “common trait currencies” in the sense of de Bello et al. (2025), representing a measure that is comparable across taxonomically and functionally disparate taxa while allowing trait-based analyses to be linked to (multi-)trophic interactions and ecosystem functioning across scales (Lavorel et al. 2013; Gravel et al. 2016; Marjakangas et al. 2022; Hanusch et al. 2026). Incorporating chemodiversity into multidimensional plant trait spaces closes a persistent Eltonian gap in trait-based ecology, extending these frameworks from describing how plants grow to capturing how they interact with the living world around them.

## Data availability

Data will be made available upon publication

## Acknowledgements

We thank Ellen Kubat and Lia Botjes for their help in data collection. The authors acknowledge funding from the German Research Foundation (DFG) FOR 3000/2 -Ecology and Evolution of Intraspecific Chemodiversity in Plants (Project number JU2856/5-2). Open Access funding enabled and organized by Projekt DEAL.

## Author contributions

RRJ initiated the study. MH, AZ, and RRJ designed the study. MH collated and processed the data. MH, RRJ, and AZ performed the analysis. MH and RRJ wrote the first draft of the manuscript. All authors contributed to manuscript revision and approved the final version.

## Bibliography

Adler LS, Irwin RE, McArt SH, Vannette RL (2021) Floral traits affecting the transmission of beneficial and pathogenic pollinator-associated microbes. Current Opinion in Insect Science 44:1–7. 10.1016/j.cois.2020.08.006

Ali JG, Agrawal AA (2012) Specialist versus generalist insect herbivores and plant defense. Trends in Plant Science 17:293–302. 10.1016/j.tplants.2012.02.006

Aristarán M, Tigas M, Merill JB (2018) Tabula technology: Knight foundation.

Becerra JX (2015) On the factors that promote the diversity of herbivorous insects and plants in tropical forests. Proceedings of the National Academy of Sciences 112:6098–6103. 10.1073/pnas.1418643112

Bellman R (1966) Dynamic Programming. Science 153:34–37. 10.1126/science.153.3731.34

Bergmann J, Weigelt A, van der Plas F, et al (2020) The fungal collaboration gradient dominates the root economics space in plants. Science Advances 6:eaba3756. 10.1126/sciadv.aba3756

Bouwmeester H, Schuurink RC, Bleeker PM, Schiestl F (2019) The role of volatiles in plant communication. The Plant Journal 100:892–907. 10.1111/tpj.14496

Bruelheide H, Dengler J, Purschke O, et al (2018) Global trait–environment relationships of plant communities. Nat Ecol Evol 2:1906–1917. 10.1038/s41559-018-0699-8

Carmona CP, Beccari E (2025) The path toward a unified trait space: synthesizing plant functional diversity. New Phytologist 248:2236–2242. 10.1111/nph.70584

Carmona CP, Bueno CG, Toussaint A, et al (2021) Fine-root traits in the global spectrum of plant form and function. Nature 597:683–687. 10.1038/s41586-021-03871-y

Chadwick S (ORCID:0009000520363751), Henderson D (ORCID:0000000335230175), Forrister DL (ORCID:0000000181707187), et al (2026) Chemical properties of foliar metabolomes represent a key axis of functional trait variation in forests of the tropical Andes. Proceedings of the Royal Society B: Biological Sciences 293:. 10.1098/rspb.2025.1721

Chiu C-H, Chao A (2014) Distance-Based Functional Diversity Measures and Their Decomposition: A Framework Based on Hill Numbers. PLOS ONE 9:e100014. 10.1371/journal.pone.0100014

Cosmo LG, Yamaguchi LF, Felix GMF, et al (2021) From the leaf to the community: Distinct dimensions of phytochemical diversity shape insect–plant interactions within and among individual plants. Journal of Ecology 109:2475–2487. 10.1111/1365-2745.13659

de Bello F, Fischer FM, Puy J, et al (2025) Raunkiæran shortfalls: Challenges and perspectives in trait-based ecology. Ecological Monographs 95:e70018. 10.1002/ecm.70018

Dehling DM, Stouffer DB (2018) Bringing the Eltonian niche into functional diversity. Oikos 127:1711–1723. 10.1111/oik.05415

Dias ATC, Rosado BHP, De Bello F, et al (2020) Alternative plant designs: consequences for community assembly and ecosystem functioning. Ann Bot 125:391–398. 10.1093/aob/mcz180

Díaz S, Kattge J, Cornelissen JHC, et al (2016) The global spectrum of plant form and function. Nature 529:167–171. 10.1038/nature16489

Dötterl S, Gershenzon J (2023) Chemistry, biosynthesis and biology of floral volatiles: roles in pollination and other functions. Nat Prod Rep 40:1901–1937. 10.1039/D3NP00024A

Eilers EJ, Kleine S, Eckert S, et al (2021) Flower Production, Headspace Volatiles, Pollen Nutrients, and Florivory in Tanacetum vulgare Chemotypes. Front Plant Sci 11:. 10.3389/fpls.2020.611877

Erb M (2019) Plant Biology: Evolution of Volatile-Mediated Plant–Plant Interactions. Current Biology 29:R873–R875. 10.1016/j.cub.2019.07.066

E-Vojtkó A, Junker RR, de Bello F, Götzenberger L (2022) Floral and reproductive traits are an independent dimension within the plant economic spectrum of temperate central Europe. New Phytologist 236:1964–1975. 10.1111/nph.18386

Fazayeli F, Banerjee A, Kattge J, et al (2014) Uncertainty Quantified Matrix Completion Using Bayesian Hierarchical Matrix Factorization. In: 2014 13th International Conference on Machine Learning and Applications. pp 312–317

Gargallo-Garriga A, Sardans J, Granda V, et al (2020) Different “metabolomic niches” of the highly diverse tree species of the French Guiana rainforests. Sci Rep 10:6937. 10.1038/s41598-020-63891-y

Gravel D, Albouy C, Thuiller W (2016) The meaning of functional trait composition of food webs for ecosystem functioning. Philos Trans R Soc Lond B Biol Sci 371:20150268. 10.1098/rstb.2015.0268

Hammerbacher A, Coutinho TA, Gershenzon J (2019) Roles of plant volatiles in defence against microbial pathogens and microbial exploitation of volatiles. Plant, Cell & Environment 42:2827–2843. 10.1111/pce.13602

Hanusch M, Dötterl S, Larue-Kontić A-AC, et al (2025) Floral scent chemodiversity is associated with high floral visitor but low bacterial richness on flowers. New Phytol 248:3270–3279. 10.1111/nph.70600

Hanusch M, Dussarrat T, Xiao X, et al (2026) Ecological role of emergent properties in the chemodiversity landscape. Nat Ecol Evol. 10.1038/s41559-026-03057-7

Jin Y, Qian H (2022) V.PhyloMaker2: An updated and enlarged R package that can generate very large phylogenies for vascular plants. Plant Diversity 44:335–339. 10.1016/j.pld.2022.05.005

Jin Y, Qian H (2023) U.PhyloMaker: An R package that can generate large phylogenetic trees for plants and animals. Plant Diversity 45:347–352. 10.1016/j.pld.2022.12.007

Junker RR (2018) A biosynthetically informed distance measure to compare secondary metabolite profiles. Chemoecology 28:29–37. 10.1007/s00049-017-0250-4

Junker RR (2016) Multifunctional and Diverse Floral Scents Mediate Biotic Interactions Embedded in Communities. In: Blande JD, Glinwood R (eds) Deciphering Chemical Language of Plant Communication. Springer International Publishing, Cham, pp 257– 282

Junker RR, Blüthgen N (2010) Floral scents repel facultative flower visitors, but attract obligate ones. Annals of Botany 105:777–782. 10.1093/aob/mcq045

Junker RR, Tholl D (2013) Volatile Organic Compound Mediated Interactions at the Plant-Microbe Interface. J Chem Ecol 39:810–825. 10.1007/s10886-013-0325-9

Kattge J, Díaz S, Lavorel S, et al (2011) TRY – a global database of plant traits. Glob Chang Biol 17:2905–2935. 10.1111/j.1365-2486.2011.02451.x

Kessler A, Baldwin IT (2001) Defensive function of herbivore-induced plant volatile emissions in nature. Science 291:2141–2144. 10.1126/science.291.5511.2141

Kessler A, Kalske A (2018) Plant Secondary Metabolite Diversity and Species Interactions. Annu Rev Ecol Evol Syst 49:115–138. 10.1146/annurev-ecolsys-110617-062406

Kindt R (2020) WorldFlora: An R package for exact and fuzzy matching of plant names against the World Flora Online taxonomic backbone data. Applications in Plant Sciences 8:e11388

Lajeunesse MJ (2016) Facilitating systematic reviews, data extraction and meta-analysis with the metagear package for r. Methods in Ecology and Evolution 7:323–330. 10.1111/2041-210X.12472

Larue A-AC, Raguso RA, Junker RR (2016) Experimental manipulation of floral scent bouquets restructures flower–visitor interactions in the field. Journal of Animal Ecology 85:396–408. 10.1111/1365-2656.12441

Lavorel S, Storkey J, Bardgett RD, et al (2013) A novel framework for linking functional diversity of plants with other trophic levels for the quantification of ecosystem services. Journal of Vegetation Science 24:942–948. 10.1111/jvs.12083

Lê S, Josse J, Husson F (2008) FactoMineR: An R Package for Multivariate Analysis. Journal of Statistical Software 25:1–18. 10.18637/jss.v025.i01

Lewinsohn TM, Novotny V, Basset Y (2005) Insects on Plants: Diversity of Herbivore Assemblages Revisited. Annual Review of Ecology, Evolution, and Systematics 36:597 –620. 10.1146/annurev.ecolsys.36.091704.175520

Loreto F, Schnitzler J-P (2010) Abiotic stresses and induced BVOCs. Trends Plant Sci 15:154 –166. 10.1016/j.tplants.2009.12.006

Marjakangas E-L, Muñoz G, Turney S, et al (2022) Trait-based inference of ecological network assembly: A conceptual framework and methodological toolbox. Ecological Monographs 92:e1502. 10.1002/ecm.1502

Marks CO, Lechowicz MJ (2006) Alternative designs and the evolution of functional diversity. Am Nat 167:55–66. 10.1086/498276

Morris EK, Caruso T, Buscot F, et al (2014) Choosing and using diversity indices: Insights for ecological applications from the German Biodiversity Exploratories. Ecology and Evolution 4:3514–3524. 10.1002/ece3.1155

Müller C, Dussarrat T, Dam N van (2026) What is a plant chemotype anyway?

Müller C, Fuchs B, Schnitzler J-P, et al (2025) Ecology and evolution of plant chemodiversity. Plant Biology 27:633–636. 10.1111/plb.70046

Müller C, Junker RR (2022) Chemical phenotype as important and dynamic niche dimension of plants. New Phytologist. 10.1111/nph.18075

Ooms J (2014) The jsonlite Package: A Practical and Consistent Mapping Between JSON Data and R Objects. arXiv:14032805 [statCO]

Paradis E, Schliep K (2019) ape 5.0: an environment for modern phylogenetics and evolutionary analyses in {R}. Bioinformatics 35:526–528

Peeters PJ, Read J, Sanson GD (2001) Variation in the guild composition of herbivorous insect assemblages among co-occurring plant species. Austral Ecology 26:385–399. 10.1046/j.1442-9993.2001.01123.x

Petrén H, Anaia RA, Aragam KS, et al (2024) Understanding the chemodiversity of plants: Quantification, variation and ecological function. Ecological Monographs e1635. 10.1002/ecm.1635

Petrén H, Köllner TG, Junker RR (2023) Quantifying chemodiversity considering biochemical and structural properties of compounds with the R package chemodiv. New Phytologist 237:2478–2492. 10.1111/nph.18685

Pinheiro J, Bates D, Team RC (2025) nlme: linear and nonlinear mixed effects models. R package version 3.1-168. The R Foundation for Statistical Computing (CRAN)

Poelen J, Gosnell S, Slyusarev S, Waters H (2026) rglobi: Interface to Global Biotic Interactions

Poelen JH, Simons JD, Mungall CJ (2014) Global biotic interactions: An open infrastructure to share and analyze species-interaction datasets. Ecological Informatics 24:148–159. 10.1016/j.ecoinf.2014.08.005

Poelman EH, van Loon JJA, Dicke M (2008) Consequences of variation in plant defense for biodiversity at higher trophic levels. Trends in Plant Science 13:534–541. 10.1016/j.tplants.2008.08.003

Raguso RA (2008) Wake up and Smell the Roses: The Ecology and Evolution of Floral Scent. Annual Review of Ecology, Evolution, and Systematics 39:549–569

Richards LA, Dyer LA, Forister ML, et al (2015) Phytochemical diversity drives plant–insect community diversity. Proceedings of the National Academy of Sciences 112:10973– 10978. 10.1073/pnas.1504977112

Richards LA, Glassmire AE, Ochsenrider KM, et al (2016) Phytochemical diversity and synergistic effects on herbivores. Phytochem Rev 15:1153–1166. 10.1007/s11101-016-9479-8

Salazar D, Lokvam J, Mesones I, et al (2018) Origin and maintenance of chemical diversity in a species-rich tropical tree lineage. Nat Ecol Evol 2:983–990. 10.1038/s41559-018-0552-0

Sandra Díaz, Jens Kattge, Johannes H. C. Cornelissen, et al (2022) The global spectrum of plant form and function: enhanced species-level trait dataset. Scientific Data 9:. 10.1038/s41597-022-01774-9

Schiestl FP, Johnson SD (2013) Pollinator-mediated evolution of floral signals. Trends in Ecology & Evolution 28:307–315. 10.1016/j.tree.2013.01.019

Sedio BE, Rojas Echeverri JC, Boya P. CA, Wright SJ (2017) Sources of variation in foliar secondary chemistry in a tropical forest tree community. Ecology 98:616–623. 10.1002/ecy.1689

Smith SA, Brown JW (2018) Constructing a broadly inclusive seed plant phylogeny. American Journal of Botany 105:302–314. 10.1002/ajb2.1019

Thon F m., Müller C, Wittmann MJ (2024) The evolution of chemodiversity in plants-From verbal to quantitative models. Ecology Letters. 10.1111/ele.14365

Viechtbauer W (2025) metafor: Meta-Analysis Package for R

Violle C, Navas M, Vile D, et al (2007) Let the concept of trait be functional! Oikos 116:882–892. 10.1111/j.0030-1299.2007.15559.x

Vlot AC, Rosenkranz M (2022) Volatile compounds—the language of all kingdoms? J Exp Bot 73:445–448. 10.1093/jxb/erab528

Volf M, Hrcek J, Julkunen-Tiitto R, Novotny V (2015a) To each its own: differential response of specialist and generalist herbivores to plant defence in willows. Journal of Animal Ecology 84:1123–1132. 10.1111/1365-2656.12349

Volf M, Julkunen-Tiitto R, Hrcek J, Novotny V (2015b) Insect herbivores drive the loss of unique chemical defense in willows. Entomologia Experimentalis et Applicata 156:88– 98. 10.1111/eea.12312

Walker TWN, Schrodt F, Allard P-M, et al (2023) Leaf metabolic traits reveal hidden dimensions of plant form and function. Sci Adv 9:eadi4029. 10.1126/sciadv.adi4029

Wan N-F, Deng J-Y, Huang K-H, et al (2017) Nucleopolyhedrovirus infection enhances plant defences by increasing plant volatile diversity. Biocontrol Science and Technology 27:1292–1307. 10.1080/09583157.2017.1393498

Wetzel WC, Whitehead SR (2020) The many dimensions of phytochemical diversity: linking theory to practice. Ecology Letters 23:16–32. 10.1111/ele.13422

Wickham H (2023) httr: Tools for Working with URLs and HTTP

William Revelle (2026) psych: Procedures for Psychological, Psychometric, and Personality Research. Northwestern University, Evanston, Illinois

Wright IJ, Reich PB, Westoby M, et al (2004) The worldwide leaf economics spectrum. Nature 428:821–827. 10.1038/nature02403

Yang H, Zhang P, Wang G, et al (2026) Positioning root chemical defence functions in the root economics space across subtropical tree species. New Phytologist n/a: 10.1111/nph.71421

Zanne AE, Tank DC, Cornwell WK, et al (2014) Three keys to the radiation of angiosperms into freezing environments. Nature 506:89–92. 10.1038/nature12872

Zimmer R (2000) Importance of Chemical Communication in Ecology. The Biological Bulletin 198:167–167. 10.2307/1542521

Zuo Z, Weraduwage SM, Huang T, Sharkey TD (2025) How volatile isoprenoids improve plant thermotolerance. Trends in Plant Science 30:1237–1250. 10.1016/j.tplants.2025.05.004

