## Supplementary Information 1 for "Chemodiversity defines a fundamental synecological dimension of plant form and function"

Supporting Information 1A: Flowchart of the literature screening and data extraction process for floral scent VOC studies.

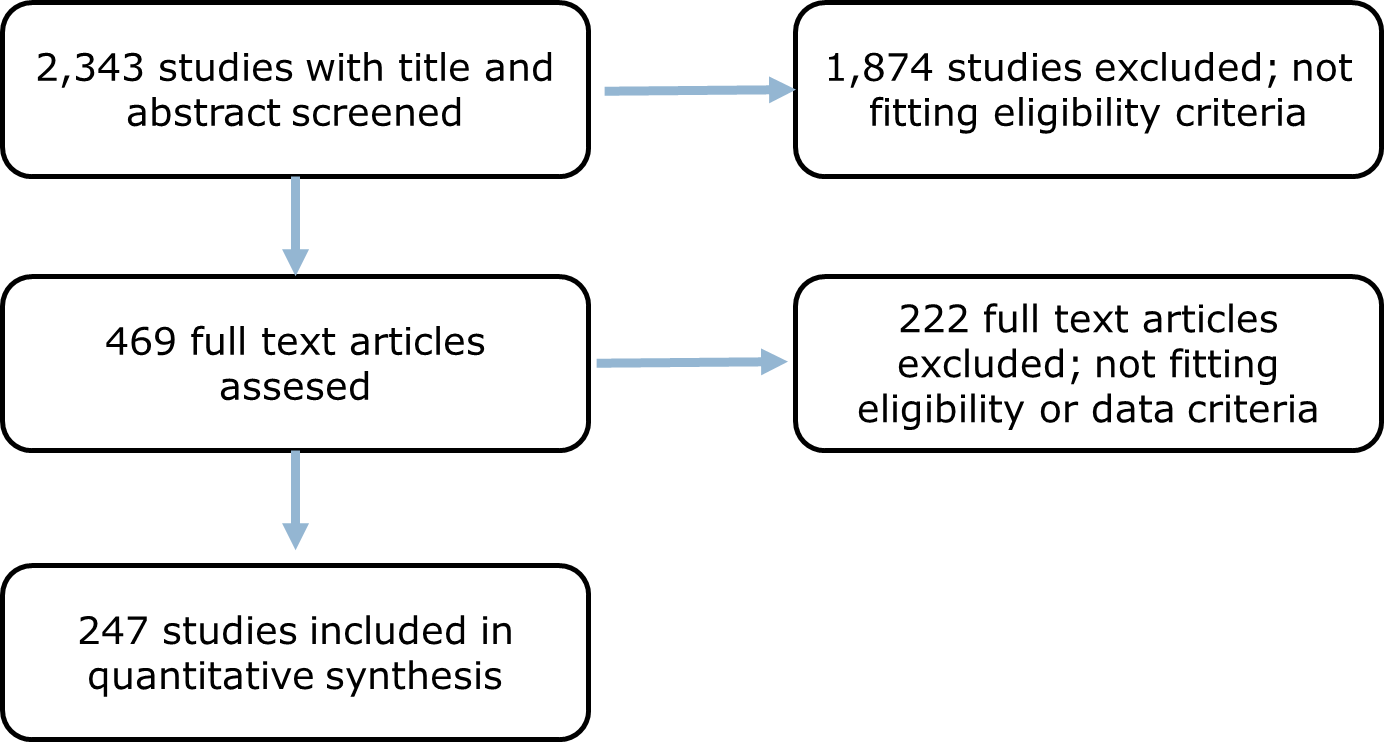

Supporting Information 1B: Flowchart of the literature screening and data extraction process for floral scent VOC studies.

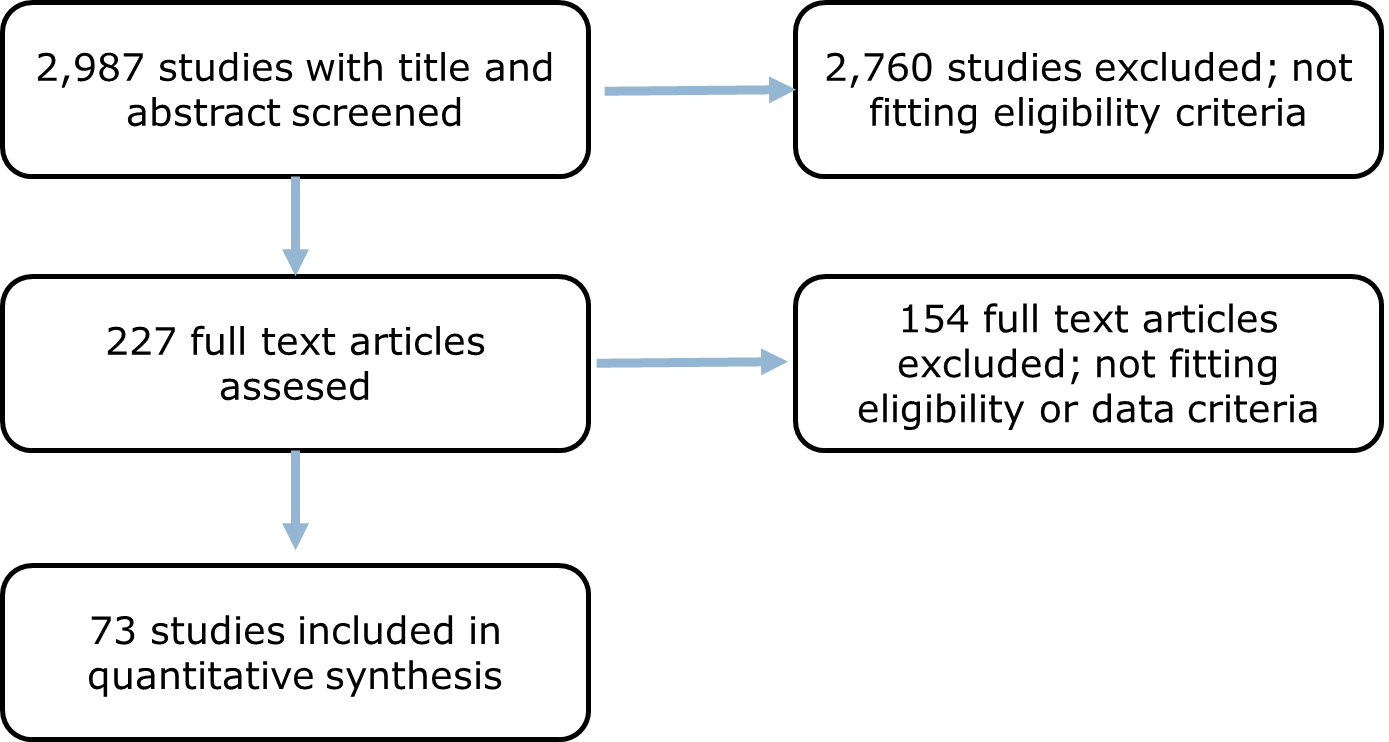
