## Supplementary Information 2 for "Chemodiversity defines a fundamental synecological dimension of plant form and function"

Supporting Information 2A: Species excluded from the final dataset after outlier screening, defined as trait values deviating by more than 4 standard deviations from the mean (|z| > 4).

| **Species** | **Affected trait(s)** |
| --- | --- |
| *Anemonella thalictroides* | Leaf Area (LA) |
| *Bonellia macrocarpa* | LMA |
| *Mauritia flexuosa* | Leaf area, LMA, Leaf Nitrogen |
| *Houttuynia cordata* | Height |
| *Trimenia moorei* | SSD |
| *Agave americana* | LMA |
| *Rhizophora stylosa* | Seed Mass |
| *Castanea crenata* | Seed Mass |
| *Eupomatia laurina* | Leaf Nitrogen |

Supporting Information 2B. Taxa removed from the interaction datasets on the family level or above because they represented implausible entries, including self-links or associated taxa that were neither pollinators nor above-ground herbivores.

Table S1: Flower dataset

| **Family** | **Reason for exclusion** |
| --- | --- |
| Araneidae | Predator |
| Bothriuridae | Predator |
| Gonyleptidae | Predator |
| Oxyopidae | Predator |
| Phytoseiidae | Predator |
| Salticidae | Predator |
| Sparassidae | Predator |
| Theridiidae | Predator |
| Thomisidae | Predator |
| Braconidae | Parasitoid; adults may take nectar but are not effective pollinators |
| Chalcididae | Parasitoid; adults may take nectar but are not effective pollinators |
| Eulophidae | Parasitoid; adults may take nectar but are not effective pollinators |
| Evaniidae | Parasitoid; adults may take nectar but are not effective pollinators |
| Gasteruptiidae | Parasitoid; adults may take nectar but are not effective pollinators |
| Ichneumonidae | Parasitoid; adults may take nectar but are not effective pollinators |
| Mymaridae | Parasitoid; adults may take nectar but are not effective pollinators |
| Platygastridae | Parasitoid; adults may take nectar but are not effective pollinators |
| Proctotrupidae | Parasitoid; adults may take nectar but are not effective pollinators |
| Pteromalidae | Parasitoid; adults may take nectar but are not effective pollinators |
| Scelionidae | Parasitoid; adults may take nectar but are not effective pollinators |
| Torymidae | Parasitoid; adults may take nectar but are not effective pollinators |
| Ixodidae | Ectoparasite of vertebrates |
| Acaridae | Detritivore |
| Philosciidae | Detritivore |
| Helicidae | Terrestrial snail |
| Agaricomycetidae | Fungal, algal or bacterial association |
| Cyclobacteriaceae | Fungal, algal or bacterial association |
| Dictyotaceae | Fungal, algal or bacterial association |
| Leptosphaeriaceae | Fungal, algal or bacterial association |
| Phaeothamniaceae | Fungal, algal or bacterial association |
| Acropomatidae | Aquatic/marine taxon |
| Cottidae | Aquatic/marine taxon |
| Hiatellidae | Aquatic/marine taxon |
| Platycephalidae | Aquatic/marine taxon |
| Pristigasteridae | Aquatic/marine taxon |
| Sabellidae | Aquatic/marine taxon |
| Corvidae | Non-nectarivorous |
| Fringillidae | Non-nectarivorous |
| Furnariidae | Non-nectarivorous |
| Iguanidae | Non-nectarivorous |
| Parulidae | Non-nectarivorous |
| Passerellidae | Non-nectarivorous |
| Passeridae | Non-nectarivorous |
| Picidae | Non-nectarivorous |
| Procyonidae | Non-nectarivorous |
| Regulidae | Non-nectarivorous |
| Sciuridae | Non-nectarivorous |
| Turdidae | Non-nectarivorous |
| Tyrannidae | Non-nectarivorous |
| Cardinalidae | Potential flower visitor in some systems, but floral scent is likely not the primary attractant for this group |
| Galagidae | Potential flower visitor in some systems, but floral scent is likely not the primary attractant for this group |
| Gekkonidae | Potential flower visitor in some systems, but floral scent is likely not the primary attractant for this group |
| Icteridae | Potential flower visitor in some systems, but floral scent is likely not the primary attractant for this group |
| Macroscelididae | Potential flower visitor in some systems, but floral scent is likely not the primary attractant for this group |
| Thraupidae | Potential flower visitor in some systems, but floral scent is likely not the primary attractant for this group |

Table S2: Leaf dataset

| **Family** | **Reason for exclusion** |
| --- | --- |
| Aegithalidae | Predator or insectivore |
| Araneidae | Predator or insectivore |
| Braconidae | Predator or insectivore |
| Canidae | Predator or insectivore |
| Chalcididae | Predator or insectivore |
| Cisticolidae | Predator or insectivore |
| Clubionidae | Predator or insectivore |
| Cuculidae | Predator or insectivore |
| Eulophidae | Predator or insectivore |
| Eurytomidae | Predator or insectivore |
| Felidae | Predator or insectivore |
| Figitidae | Predator or insectivore |
| Furnariidae | Predator or insectivore |
| Galagidae | Predator or insectivore |
| Gekkonidae | Predator or insectivore |
| Ichneumonidae | Predator or insectivore |
| Leucospidae | Predator or insectivore |
| Macroscelididae | Predator or insectivore |
| Momotidae | Predator or insectivore |
| Muscicapidae | Predator or insectivore |
| Myrmecophagidae | Predator or insectivore |
| Oxyopidae | Predator or insectivore |
| Paridae | Predator or insectivore |
| Parulidae | Predator or insectivore |
| Picidae | Predator or insectivore |
| Platygastridae | Predator or insectivore |
| Proctotrupidae | Predator or insectivore |
| Regulidae | Predator or insectivore |
| Salticidae | Predator or insectivore |
| Sapygidae | Predator or insectivore |
| Scelionidae | Predator or insectivore |
| Scotocercidae | Predator or insectivore |
| Segestriidae | Predator or insectivore |
| Sittidae | Predator or insectivore |
| Sparassidae | Predator or insectivore |
| Sylviidae | Predator or insectivore |
| Thamnophilidae | Predator or insectivore |
| Thomisidae | Predator or insectivore |
| Torymidae | Predator or insectivore |
| Trombidiidae | Predator or insectivore |
| Vireonidae | Predator or insectivore |
| Bucerotidae | Frugivore/granivore |
| Calcariidae | Frugivore/granivore |
| Callitrichidae | Frugivore/granivore |
| Campephagidae | Frugivore/granivore |
| Capitonidae | Frugivore/granivore |
| Cardinalidae | Frugivore/granivore |
| Cebidae | Frugivore/granivore |
| Columbidae | Frugivore/granivore |
| Corvidae | Frugivore/granivore |
| Cotingidae | Frugivore/granivore |
| Emberizidae | Frugivore/granivore |
| Fringillidae | Frugivore/granivore |
| Gliridae | Frugivore/granivore |
| Icteridae | Frugivore/granivore |
| Leiothrichidae | Frugivore/granivore |
| Mitrospingidae | Frugivore/granivore |
| Muridae | Frugivore/granivore |
| Oriolidae | Frugivore/granivore |
| Passerellidae | Frugivore/granivore |
| Passeridae | Frugivore/granivore |
| Pipridae | Frugivore/granivore |
| Pitheciidae | Frugivore/granivore |
| Procyonidae | Frugivore/granivore |
| Prunellidae | Frugivore/granivore |
| Rallidae | Frugivore/granivore |
| Ramphastidae | Frugivore/granivore |
| Sciuridae | Frugivore/granivore |
| Semnornithidae | Frugivore/granivore |
| Sturnidae | Frugivore/granivore |
| Thraupidae | Frugivore/granivore |
| Tityridae | Frugivore/granivore |
| Trogonidae | Frugivore/granivore |
| Tupaiidae | Frugivore/granivore |
| Turdidae | Frugivore/granivore |
| Agaricomycetidae | Fungal association (pathogen, endophyte or saprotroph) |
| Auriculariaceae | Fungal association (pathogen, endophyte or saprotroph) |
| Botryosphaeriaceae | Fungal association (pathogen, endophyte or saprotroph) |
| Chaetomiaceae | Fungal association (pathogen, endophyte or saprotroph) |
| Clavariaceae | Fungal association (pathogen, endophyte or saprotroph) |
| Corticiaceae | Fungal association (pathogen, endophyte or saprotroph) |
| Cystostereaceae | Fungal association (pathogen, endophyte or saprotroph) |
| Diaporthaceae | Fungal association (pathogen, endophyte or saprotroph) |
| Dothideomycetidae | Fungal association (pathogen, endophyte or saprotroph) |
| Ganodermataceae | Fungal association (pathogen, endophyte or saprotroph) |
| Hypocreaceae | Fungal association (pathogen, endophyte or saprotroph) |
| Lachnocladiaceae | Fungal association (pathogen, endophyte or saprotroph) |
| Leotiaceae | Fungal association (pathogen, endophyte or saprotroph) |
| Massarinaceae | Fungal association (pathogen, endophyte or saprotroph) |
| Melogrammataceae | Fungal association (pathogen, endophyte or saprotroph) |
| Morchellaceae | Fungal association (pathogen, endophyte or saprotroph) |
| Mycosphaerellaceae | Fungal association (pathogen, endophyte or saprotroph) |
| Myxotrichaceae | Fungal association (pathogen, endophyte or saprotroph) |
| Nectriaceae | Fungal association (pathogen, endophyte or saprotroph) |
| Ophiostomataceae | Fungal association (pathogen, endophyte or saprotroph) |
| Patellariaceae | Fungal association (pathogen, endophyte or saprotroph) |
| Pestalotiopsidaceae | Fungal association (pathogen, endophyte or saprotroph) |
| Pezizaceae | Fungal association (pathogen, endophyte or saprotroph) |
| Pleurotaceae | Fungal association (pathogen, endophyte or saprotroph) |
| Psathyrellaceae | Fungal association (pathogen, endophyte or saprotroph) |
| Schizophyllaceae | Fungal association (pathogen, endophyte or saprotroph) |
| Sordariomycetidae | Fungal association (pathogen, endophyte or saprotroph) |
| Stictidaceae | Fungal association (pathogen, endophyte or saprotroph) |
| Thelephoraceae | Fungal association (pathogen, endophyte or saprotroph) |
| Tremellaceae | Fungal association (pathogen, endophyte or saprotroph) |
| Tricholomataceae | Fungal association (pathogen, endophyte or saprotroph) |
| Valsaceae | Fungal association (pathogen, endophyte or saprotroph) |
| Venturiaceae | Fungal association (pathogen, endophyte or saprotroph) |
| Xylariaceae | Fungal association (pathogen, endophyte or saprotroph) |
| Xylariomycetidae | Fungal association (pathogen, endophyte or saprotroph) |
| Armadillidiidae | Detritivore/decomposer or aquatic larval stage |
| Blaniulidae | Detritivore/decomposer or aquatic larval stage |
| Chironomidae | Detritivore/decomposer or aquatic larval stage |
| Culicidae | Detritivore/decomposer or aquatic larval stage |
| Ellobiidae | Detritivore/decomposer or aquatic larval stage |
| Ephydridae | Detritivore/decomposer or aquatic larval stage |
| Gastrodontidae | Detritivore/decomposer or aquatic larval stage |
| Glomeridae | Detritivore/decomposer or aquatic larval stage |
| Nemasomatidae | Detritivore/decomposer or aquatic larval stage |
| Porcellionidae | Detritivore/decomposer or aquatic larval stage |
| Sciaridae | Detritivore/decomposer or aquatic larval stage |
| Simuliidae | Detritivore/decomposer or aquatic larval stage |
| Stratiomyidae | Detritivore/decomposer or aquatic larval stage |
| Tipulidae | Detritivore/decomposer or aquatic larval stage |
| Vitrinidae | Detritivore/decomposer or aquatic larval stage |
| Acropomatidae | Aquatic/marine taxon |
| Cheloniidae | Aquatic/marine taxon |
| Cottidae | Aquatic/marine taxon |
| Cyprinidae | Aquatic/marine taxon |
| Lampridae | Aquatic/marine taxon |
| Mytilidae | Aquatic/marine taxon |
| Serrasalmidae | Aquatic/marine taxon |
| Phyllostomidae | Nectarivore/frugivore |
| Zosteropidae | Nectarivore/frugivore |
| Agriolimacidae | Herbivore, but likely contact chemoreception, not olfaction |
| Helicidae | Herbivore, but likely contact chemoreception, not olfaction |
| Veronicellidae | Herbivore, but likely contact chemoreception, not olfaction |
| Bovidae | Herbivore, but generalist with non-olfactory host selection |
| Camelidae | Herbivore, but generalist with non-olfactory host selection |
| Equidae | Herbivore, but generalist with non-olfactory host selection |
| Elephantidae | Herbivore, but generalist with non-olfactory host selection |
| Tapiridae | Herbivore, but generalist with non-olfactory host selection |
| Leporidae | Herbivore, but generalist with non-olfactory host selection |
| Myocastoridae | Herbivore, but likely visual or gustatory foraging |
| Bradypodidae | Herbivore, but likely visual or gustatory foraging |
| Atelidae | Herbivore, but likely visual or gustatory foraging |
| Cercopithecidae | Herbivore, but likely visual or gustatory foraging |
| Hominidae | Herbivore, but likely visual or gustatory foraging |
| Cracidae | Herbivore, but likely visual or gustatory foraging |
| Emydidae | Herbivore, but likely visual or gustatory foraging |
| Iguanidae | Herbivore, but likely visual or gustatory foraging |
| Testudinidae | Herbivore, but likely visual or gustatory foraging |
| Suidae | Omnivore |
