## Supplementary Information 3 for "Chemodiversity defines a fundamental synecological dimension of plant form and function"

**Table S3A.** Output of the exploratory factor analysis of six plant economic spectrum traits (leaf area, nitrogen content, leaf mass per area, plant height, seed mass, and stem specific density) and log-transformed floral VOC chemodiversity (maximum likelihood estimation, promax rotation). Standardized factor loadings (ML1, ML2) with bootstrapped 95% confidence intervals are shown, together with communalities (h²), uniqueness (u²), item complexity (com), and model fit statistics.

| Section | Item | ML1 | ML1_CI | ML2 | ML2_CI | h2 | u2 | com |
| --- | --- | --- | --- | --- | --- | --- | --- | --- |
| **Factor loadings** | LA | **0.620** | [0.53, 0.71] | -0.28 | [-0.41, -0.16] | 0.35 | 0.65 | 1.38 |
|  | Nitrogen | 0.120 | [0.05, 0.19] | **-0.67** | [-0.79, -0.56] | 0.41 | 0.59 | 1.06 |
|  | LMA | 0.040 | [-0.02, 0.12] | **0.81** | [0.69, 0.95] | 0.68 | 0.32 | 1.00 |
|  | Height | **0.980** | [0.89, 1.03] | 0.03 | [-0.04, 0.11] | 0.97 | 0.03 | 1.00 |
|  | SeedMass | **0.650** | [0.57, 0.73] | 0.09 | [0.01, 0.18] | 0.47 | 0.53 | 1.04 |
|  | SSD | **0.530** | [0.46, 0.59] | **0.50** | [0.42, 0.61] | 0.71 | 0.29 | 2.00 |
|  | Chemodiversity (flower) | -0.180 | [-0.29, -0.06] | -0.03 | [-0.16, 0.1] | 0.04 | 0.96 | 1.05 |
| **Model fit** | RMSR | 0.036 |  |  |  |  |  |  |
|  | TLI | 0.928 |  |  |  |  |  |  |
|  | RMSEA | 0.092 |  |  |  |  |  |  |
|  | RMSEA lower 90% CI | 0.060 |  |  |  |  |  |  |
|  | RMSEA upper 90% CI | 0.127 |  |  |  |  |  |  |
|  | BIC | -15.043 |  |  |  |  |  |  |
|  | Chi-square | 31.844 |  |  |  |  |  |  |
|  | df | 8.000 |  |  |  |  |  |  |
|  | p-value | 0.000 |  |  |  |  |  |  |

**Table S3B.** Output of the exploratory factor analysis of six plant economic spectrum traits (leaf area, nitrogen content, leaf mass per area, plant height, seed mass, and stem specific density) and log-transformed foliar VOC chemodiversity (maximum likelihood estimation, promax rotation). Standardized factor loadings (ML1, ML2) with bootstrapped 95% confidence intervals are shown, together with communalities (h²), uniqueness (u²), item complexity (com), and model fit statistics.

| Section | Item | ML1 | ML1_CI | ML2 | ML2_CI | h2 | u2 | com |
| --- | --- | --- | --- | --- | --- | --- | --- | --- |
| **Factor loadings** | LA | **0.560** | [0.43, 0.68] | **-0.44** | [-0.61, -0.24] | 0.40 | 0.60 | 1.89 |
|  | Nitrogen | -0.060 | [-0.19, 0.08] | **-0.66** | [-0.85, -0.51] | 0.46 | 0.54 | 1.01 |
|  | LMA | 0.040 | [-0.07, 0.18] | **0.92** | [0.72, 1.08] | 0.86 | 0.14 | 1.00 |
|  | Height | **0.810** | [0.7, 0.94] | 0.27 | [0.15, 0.4] | 0.82 | 0.18 | 1.23 |
|  | SeedMass | **0.720** | [0.61, 0.84] | 0.00 | [-0.12, 0.13] | 0.52 | 0.48 | 1.00 |
|  | SSD | **0.690** | [0.58, 0.82] | 0.36 | [0.26, 0.48] | 0.71 | 0.29 | 1.52 |
|  | Chemodiversity (leaf) | -0.020 | [-0.23, 0.18] | -0.26 | [-0.49, -0.04] | 0.07 | 0.93 | 1.01 |
| **Model fit** | RMSR | 0.034 |  |  |  |  |  |  |
|  | TLI | 0.992 |  |  |  |  |  |  |
|  | RMSEA | 0.029 |  |  |  |  |  |  |
|  | RMSEA lower 90% CI | 0.000 |  |  |  |  |  |  |
|  | RMSEA upper 90% CI | 0.121 |  |  |  |  |  |  |
|  | BIC | -28.577 |  |  |  |  |  |  |
|  | Chi-square | 8.806 |  |  |  |  |  |  |
|  | df | 8.000 |  |  |  |  |  |  |
|  | p-value | 0.359 |  |  |  |  |  |  |
